# Eco-evolutionary games in noisy environments

**DOI:** 10.64898/2026.05.20.726658

**Authors:** Fantine Bodin, Guocheng Wang, Joshua B. Plotkin

## Abstract

Cooperative and competitive interactions among individuals harvesting resources can shape environmental states, such as prey abundance. In turn, environmental conditions feed back to influence strategic interactions. Eco-evolutionary game theory studies how these feedbacks shape the co-evolution of behavior and environment. Existing models typically assume deterministic, noise-free environmental dynamics. However, real environments are inherently stochastic, for example due to finite resources, and noise can qualitatively alter social outcomes. Here, we incorporate stochastic environmental dynamics into eco-evolutionary game theory. When environmental change is slow relative to strategy updates, we show that behavior reflects a mixture of the games associated with low and high environmental states, often yielding outcomes qualitatively distinct from deterministic predictions. In particular, environmental stochasticity can eliminate bistability and enforce dominance of a single behavior. When environmental dynamics are faster, populations have less opportunity to track fluctuations, and behavior converges toward strategies that are optimal on average. Stochasticity can even causes persistent oscillations in the tragedy of commons, in regimes where classical models predict stability. Our framework provides a tractable approach for analyzing social behavior linked to environmental dynamics how noise shapes long-term eco-evolutionary outcomes.

## 1 Introduction

Evolutionary game theory describes behavioral dynamics in a population of interacting individuals who have limited capacity for cognition and whose behavior are determined by fitness. Recently, this field has been extended to accommodate strategic interactions that are mediated by the state of the environment [20, 27, 11]. In many settings, individual’s actions directly affect how environmental resources are consumed, transformed, or produced [7]. As a result, individual behavior feeds back onto the dynamics of the environment itself. For example, humans may choose to harvest fish intensively (a high-impact, defection-like behavior), or to restrain harvesting in order to preserve the resource and ensure its long-term availability (a low-impact, cooperative behavior). Conversely, the state of the environment influences the benefits and costs associated with different behaviors, thereby shaping the evolutionary dynamics of strategies in the population. For instance, cooperation may be more appealing to individuals when fish stocks are depleted, whereas selfish harvesting may be favored when resources are abundant. These reciprocal interactions between behavior and environmental state are known as eco-evolutionary feedbacks and form the basis of eco-evolutionary game theory (Fig. 1).

**Figure 1:**
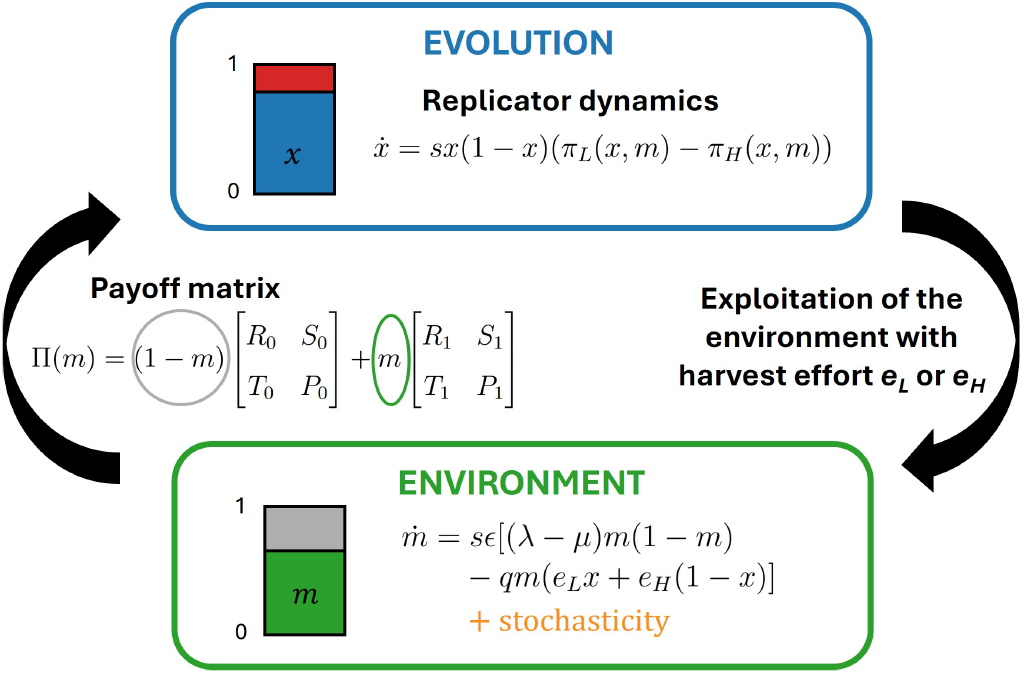
Schematic of eco-evolutionary games. The variable *x* denotes the proportion of cooperators in the population, and *m* denotes the level of the environmental resource. The resource *m* (e.g., the stock of fish available for harvest) follows intrinsic logistic dynamics governed by a birth rate *λ* and a death rate *µ*. Cooperators exert a low impact on the environment (harvest effort *e*_*L*_), whereas defectors exert a higher impact (harvest effort *e*_*H*_, with *e*_*L*_ *< e*_*H*_); thus, the population composition *x* modifies the dynamics of *m*. The environmental state *m*, in turn, determines the payoff structure of social interactions: individuals play one game when resources are depleted and another when resources are abundant. Strategy frequencies evolve through payoff-biased imitation, captured by a replicator-type dynamic. Because strategy frequencies affect the environment and the environment feeds back to shape payoffs, the system exhibits ecoevolutionary feedbacks. The parameter *s* denotes the strength of selection, and *ϵ* quantifies the timescale of environmental change relative to strategy evolution. All the variables are summarized in Table 1 in Supplementary Information. Unlike earlier deterministic models of eco-evolutionary game theory [20], we incorporate stochasticity into the environmental dynamics (shown in orange).

[20] provided a systematic overview of the possible dynamical outcomes of such eco-evolutionary systems. They showed that coupled strategy–environment dynamics are governed by the incentives to lead or to follow strategic change within the population. [11] provided a more systematic approach to understanding these outcomes by considering class of social dilemma (game) played under depleted versus replete environmental conditions (prisoner’s dilemma, stag-hunt, snowdrift, or harmony game). One particularly striking outcome occurs when individuals have incentives both to lead and to follow a “gold rush” toward resource exploitation, as well as incentives to lead and follow an environmental movement towards conservation: this occurs when the effective game is a harmony game in the resource-poor environmental state and a prisoner’s dilemma in the resource-rich environmental state. The resulting coupled dynamics can exhibit persistent cycles [21], which have been called an oscillatory tragedy of the commons by [27**?**].

Early studies in eco-evolutionary game theory assumed that environmental change is driven solely by the actions of the population [27, 11]. [20] introduced a framework in which resources also possess intrinsic dynamics, in addition to external modification by the actions of individuals. In their model, resources may be intrinsically renewing (following logistic growth), decaying, or governed by tipping points. In all of these studies, however, environmental dynamics are treated as strictly deterministic.

Stochasticity in environmental resources is commonplace, in reality, and it may have important consequences for the evolution of strategies in a population (Fig. 1). Indeed, it is well known, outside the context of eco-evolutionary games, that noise can qualitatively alter evolutionary dynamic. For example, demographic noise in the population of players can sustain a positive frequency of cooperation in populations that would otherwise be dominated by defection without noise [5, 24]. In classical evolutionary games without environmental feedbacks, noise in payoff observations [22] or in social environments [23] can also produce qualitative changes in behavioral outcomes. When payoffs fluctuate over time and differ across interaction types, novel dynamics can emerge, such as stable coexistence of both cooperation and defection in the prisoner’s dilemma. This raises the natural question of how behavior will evolve when payoffs depend on a coupled environmental state that is also subject to stochastic fluctuations.

Here, we extend eco-evolutionary game theory by explicitly incorporating stochasticity in the environmental dynamics. We examine how environmental noise affects the evolution of strategies in a population, asking whether stochasticity induces merely quantitative shifts or fundamentally qualitative changes in behavioral outcomes; how it matters compared with deterministic dynamics; and how outcomes depend on the magnitude of environmental fluctuations. We focus on an environment whose intrinsic dynamics are governed by a birth–death process with density-dependent mortality. This process is inherently stochastic: resource abundance fluctuates around the environment’s carrying capacity. When the carrying capacity is large, these dynamics closely approximate deterministic logistic growth. By contrast, when the carrying capacity is small, environmental dynamics become highly noisy—so-called demographic noise—and resource levels more frequently approach very low values, potentially leading to resource extinction in the absence of immigration.

A natural example of real-world eco-evolutionary dynamics with noise is a population of individuals who harvest fish from a lake with a small carrying capacity, for example due to strong competition among fishes. In such a system, fish (resource) abundance will be subject to substantial stochastic fluctuations. What are the implications of this environmental noise for the behavior of harvesters and long-term dynamics of the resource?

This study is organized as follows. First we derive the form of environmental demographic noise from a microscopic, individual-based model; and we incorporate these stochastic dynamics into the eco-evolutionary framework. We then analyze the evolution of behavior under the assumption that environmental change is slow relative to strategy updating. In this regime, we show that environmental noise can qualitatively alter long-term outcomes: population behavior becomes a structured mixture of the games played at different environmental resource levels, and stochasticity can eliminate bistability that would arise in deterministic settings. We then consider more generally how the environmental carrying capacity (and, hence, strength of stochasticity) and the timescale of environmental change impact eco-evolutionary dynamics. We close by focusing on the special case of an oscillatory tragedy of commons—where we find that oscillations persist in the presence of noise even in regimes where they would be otherwise be extinguished.

## 2 Model

### 2.1 Model of strategy evolution

We consider a population of *N* individuals who interact. Each individual adopts one of two strategies for a social interaction: a low-impact strategy (cooperation, denoted L) or a high-impact strategy (defection, denoted H), and their immediate payoffs depend upon what strategies they and their co-player employ. Over time, individuals update their strategies by imitating others with higher payoff [15], following a standard payoff-biased imitation framework [19, 12]. We consider an imitation process in which ordered pairs of individuals *i* and *j* are chosen uniformly for a potential strategy update. Individual *i* then copies the strategy of *j* with a probability that depends on their two finesses. The fitness of each individual depends on their strategy and the frequency of strategies in the population: we denote by *π*_*L*_ the fitness of the low-impact strategy (summed across all pairwise games); and by *π*_*H*_ the fitness of the high-impact strategy. The probability of *i* copying *j* is then *ϕ*(*π*_*i*_, *π*_*j*_) = 1*/*(1 + exp[*s*(*π*_*i*_ − *π*_*j*_)], where the parameter *s* is the strength of selection, which is assumed to be weak (*s* ≪ 1) [**?**]. The fraction of cooperators *x* in the population then follows a discrete Markov Chain process in continuous time with the following events:

- *x* increases from *x* to *x*+1*/N* at rate (1−*x*)*x ϕ*(*π*_*H*_, *π*_*L*_), corresponding to a defector imitating a cooperator.
- *x* decreases from *x* to *x* − 1*/N* at rate *x*(1 − *x*) *ϕ*(*π*_*L*_, *π*_*H*_), corresponding to a cooperator imitating a defector.

### 2.2 Model of environmental dynamics

The dynamics of the environment are governed by an intrinsic growth as well as external perturbations caused by the actions of the individuals. We model the intrinsic dynamics of the environmental resource as a discrete birth-death process, with density-dependent death rates, in continuous time. Environmental resources, for example fish stock, are discrete and give birth at a rate *λ* and die at a rate *µ*. Density-dependence regulation, due to competition between individuals, stabilizes the resource abundance around a carrying capacity *K*.

Let *M* ∈ ℕ be the number of environmental resources. The intrinsic dynamics of *M* are described by the following continuous-time Markov process [6]:

- *M* increases from *M* to *M* + 1 at rate λ*M*.
- *M* decreases from *M* to *M* − 1 at rate 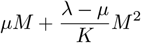.

Here *λ* − *µ* represents the net growth rate per capita, and 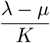 quantifies the strength of density-dependent competition.

We next incorporate the effects of individual behavior on resource dynamics. We assume that the resource stock *M* is reduced by harvesting or consumption by individuals in the population. Cooperators employ a low-impact strategy and harvest resources at effort *e*_*L*_, whereas defectors employ a high-impact strategy and harvest at effort *e*_*H*_ *> e*_*L*_. We introduce a parameter *q* that converts aggregate harvesting pressure into a rate of resource depletion, and we assume *e*_*H*_ *< r/q* so that the resource remains positive at steady state.

In our model, individual actions affect resource births and deaths symmetrically, so that their net effect on the environment is deterministic and independent of population size. Given a fraction *x* of cooperators in the population, the resource dynamics for *M* are as follows:

- *M* increases from *M* to *M* + 1 at rate 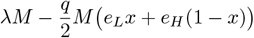
- *M* decreases from *M* to *M* − 1 at rate 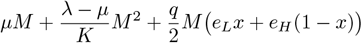.

Finally, we rescale resource abundance by the carrying capacity *K*, defining *m* = *M/K*. In the absence of harvesting, *m* stabilizes around 1. We also introduce a timescale parameter *ϵ* to account for the possibility that environmental dynamics—driven by both intrinsic growth and harvesting—occur on a different timescale than strategy evolution. Under these scalings, the dynamics of *m* are described by the following continuous-time Markov process:

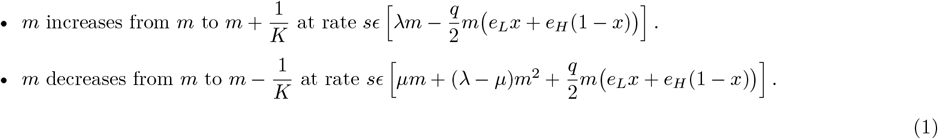

### 2.3 Model of joint eco-evolutionary dynamics

So far, we have specified how strategies evolve as a function of payoffs and how the environment evolves given the composition of strategies in the population. The environmental state, in turn, feeds back to influence the payoffs of cooperation and defection, so that the payoff matrix depends on the level of environmental resources.

We consider a linear 2 × 2 symmetric eco-evolutionary game in which the payoffs of both strategies depend on the environmental state. Individuals play game 0, denoted Π(0), when the environment is poor, and game 1, denoted Π(1), when the environment is rich. Because the environmental state is normalized by the carrying capacity *K*, it typically lies between 0 and 1; accordingly, we assume that individuals play Π(0) when *m* = 0 and Π(1) when *m* = 1.

In contrast to previous deterministic models, stochasticity in our framework allows *m* to exceed 1. When *m >* 1, we assume that individuals play the same game as when *m* = 1. Defining 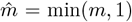 matrix as a function of the environmental state *m* is given by

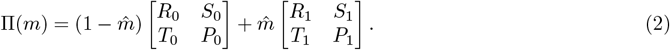

Note that in [20], because the environment is deterministic, it varies only between two equilibrium values: a value 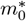, corresponding to the poorest possible environment which occurs when all individuals adopt the high-impact strategy, and a value 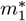, corresponding to the richest possible environment when all individuals adopt the low-impact strategy. In that framework, the environmental state was rescaled between these two values, and individuals are assumed to play Π(0) when 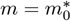 and Π(1) when 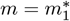.

In our model, by contrast, intrinsic stochasticity allows the environmental state *m* to fluctuate beyond these equilibrium values, reaching very low levels when high-impact strategies dominate and approaching the carrying capacity—and even exceeding it—when low-impact strategies dominate. This requires a revised definition of what constitute the poorest and richest environmental states. For simplicity, we assume that the type of game played when 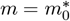 (respectively, 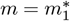) is the same as the type of game played when *m=0]* (respectively, *m=1*). We choose parameters such that 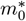 lies sufficiently close to 0 and 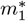 lies sufficiently close to 1.

Given these considerations, we characterize the game structure as follows, using the notation introduced in [20] (see also Table 1 in Supplementary Information):

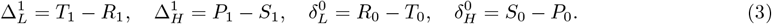

We assume that cooperation is always beneficial both to the focal individual and to its interaction partner, so that *R > P*, *R > S*, and *T > P*. Under this assumption, the qualitative structure of the game defined by each payoff matrix Π(0) and Π(1) can be classified into the following categories: *T > R* and *P > S* corresponds to a prisoner’s dilemma; *T > R* and *S > P* corresponds to a snowdrift game (also known as the chicken or hawk–dove game); *R > T* and *P > S* corresponds to a stag-hunt game (also known as a coordination game); and *R > T* and *S > P* corresponds to a harmony game (also known as trivial game).

Using Π(*m*), we write the payoffs associated with the low-impact and high-impact strategies as *π*_*L*_(*x, m*) and *π*_*H*_ (*x, m*), respectively:

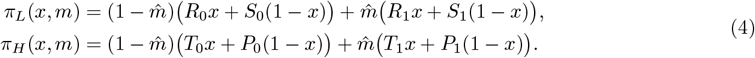

Together, equations (1) and (4) define a model of joint eco-evolutionary dynamics as two coupled stochastic processes: environmental dynamics *m* depend on the composition of strategies in the population, while the evolution of strategy frequencies depends on the environmental state through the payoff matrix Π(*m*).

### 2.4 Derivation of associated SDEs

Whereas [20] studied eco-evolutionary dynamics by analyzing the associated ordinary differential equations (*i*.*e*. replicator dynamics), our model requires an analysis based on stochastic differential equations. To derive this continuous formulation, we assume that the population size and the environmental carrying capacity are sufficiently large so that the underlying discrete eco-evolutionary process can be approximated by a continuous-state model.

We first derive an ordinary differential equation (ODE) for the proportion of cooperators by taking the limit of large population size *N*. The expected change in *x* over an update event is

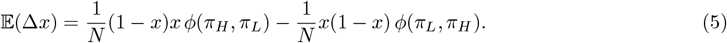

Under the common assumption of weak selection (*s* ≪ 1) [18], we expand *ϕ*(*π*_*i*_, *π*_*j*_) =(1 +exp[*s*(*π*_*i*_ − *π*_*j*_)])^*−*1^ to first order in *s*. After rescaling time, we obtain

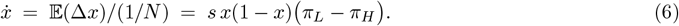

Finally, we derive a continuous model for environmental dynamics in the limit of large carrying capacity *K*, while retaining demographic noise. Rescaling time by *K* and *ϵ* by *K/N* yields a stochastic differential equation (SDE) for the environmental state *m* [6]:

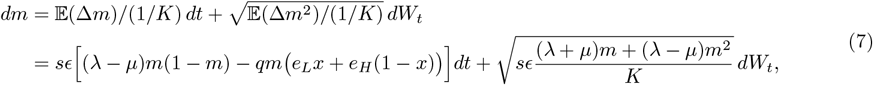

where *dW*_*t*_ denotes standard Brownian motion. Details are provided in Supplementary Information. For our analysis and our simulations, we assume that 0 is not an absorbing state but a reflecting boundary for *m*. This hypothesis reflects a low rate of immigration of environmental resources from outside the system, which prevents the resources from definitive extinction.

Note that the noise term is scaled by 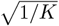. As a result, when the environmental resources have a very large carrying capacity *K*, their dynamics becomes deterministic and logistic as in [20]. If *K* is sufficiently large to consider a differential equation but not sufficiently large for the noise term to vanish, the dynamics of the environment are stochastic and this implies that the evolution of strategies is also stochastic. To sum up, we study a dynamical system with an ODE joined to an SDE:

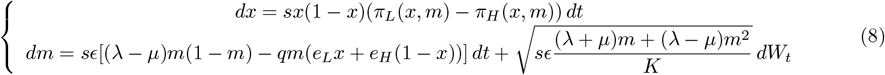

### 2.5 Timescale separation

#### Existence and stability of equilibria of *x* as a function of *m*: bifurcation diagrams

When *ϵ* is sufficiently small, environmental dynamics are slow relative to the evolution of strategies in the population. In this regime, the population effectively reaches its strategic equilibrium before the environmental state changes appreciably, yielding a separation of timescales.

For a fixed environmental state *m*, the fraction of cooperators *x* admits three equilibria:

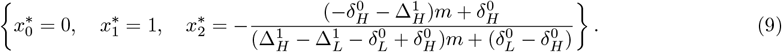

We analyze the stability of these equilibria as functions of *m* and construct bifurcation diagrams of *x*\* versus *m* for all combinations of 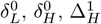, and 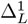 (Fig. 3). Details are provided in Supplementary Information.

**Figure 2:**
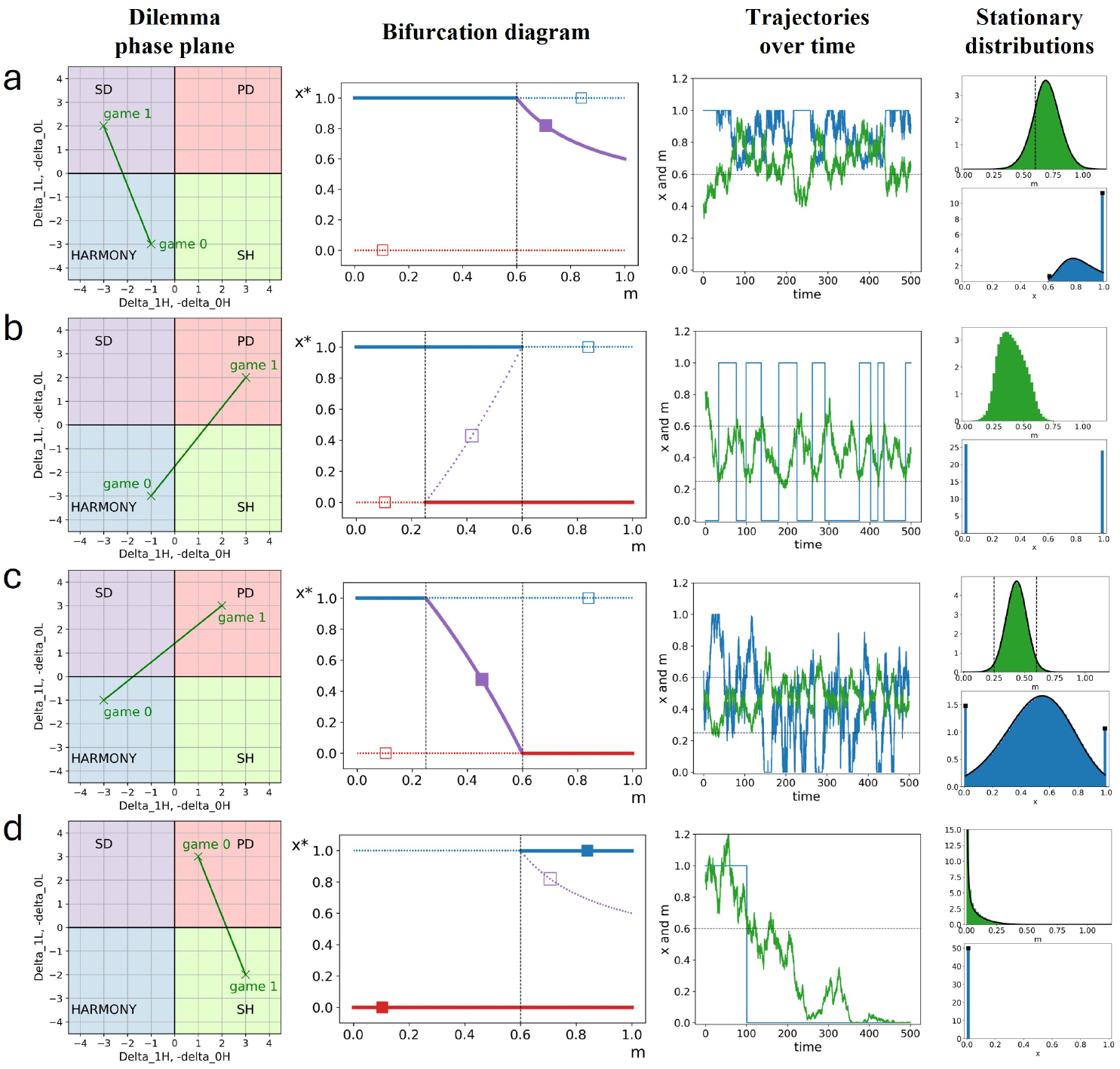
Dynamics of cooperator frequency *x* and environmental level *m* for four combinations of game types (rows). The first column illustrates the combination of games in a **dilemma phase plane** [11]: crosses denote the games played at *m* = 0 and *m* = 1, and the green line shows the effective game for intermediate environmental states. The second column shows the **bifurcation diagram** of *x*\* (the equilibrium cooperator frequency) as a function of *m*. Three types of equilibria are possible: *x* = 0 (red), *x* = 1 (blue), and an interior equilibrium *x* ∈ (0, 1) (purple), whose existence and stability depend on *m*. Solid lines indicate stable equilibria and dotted lines unstable equilibria. Squares denote the equilibria (*m, x*) without noise from [20]: filled squares are stable and open squares unstable. The third column shows representative trajectories of the environmental state *m* (green) and the corresponding cooperator frequency *x* (blue). The fourth and fifth columns display the **stationary distributions** of *m* (green) and *x* (blue), obtained from simulations and analytical calculations (black curves indicate the analytical stationary distributions, see Eqs. (**??**) and (**??**) in Supplementary Information). In row **b**, the stationary distribution cannot be computed analytically because hysteresis in the bifurcation diagram prevents *x* from being expressed as a single-valued function of *m*. All panels show dynamics under timescale separation (*ϵ* → 0) with *K* = 100.

**Figure 3:**
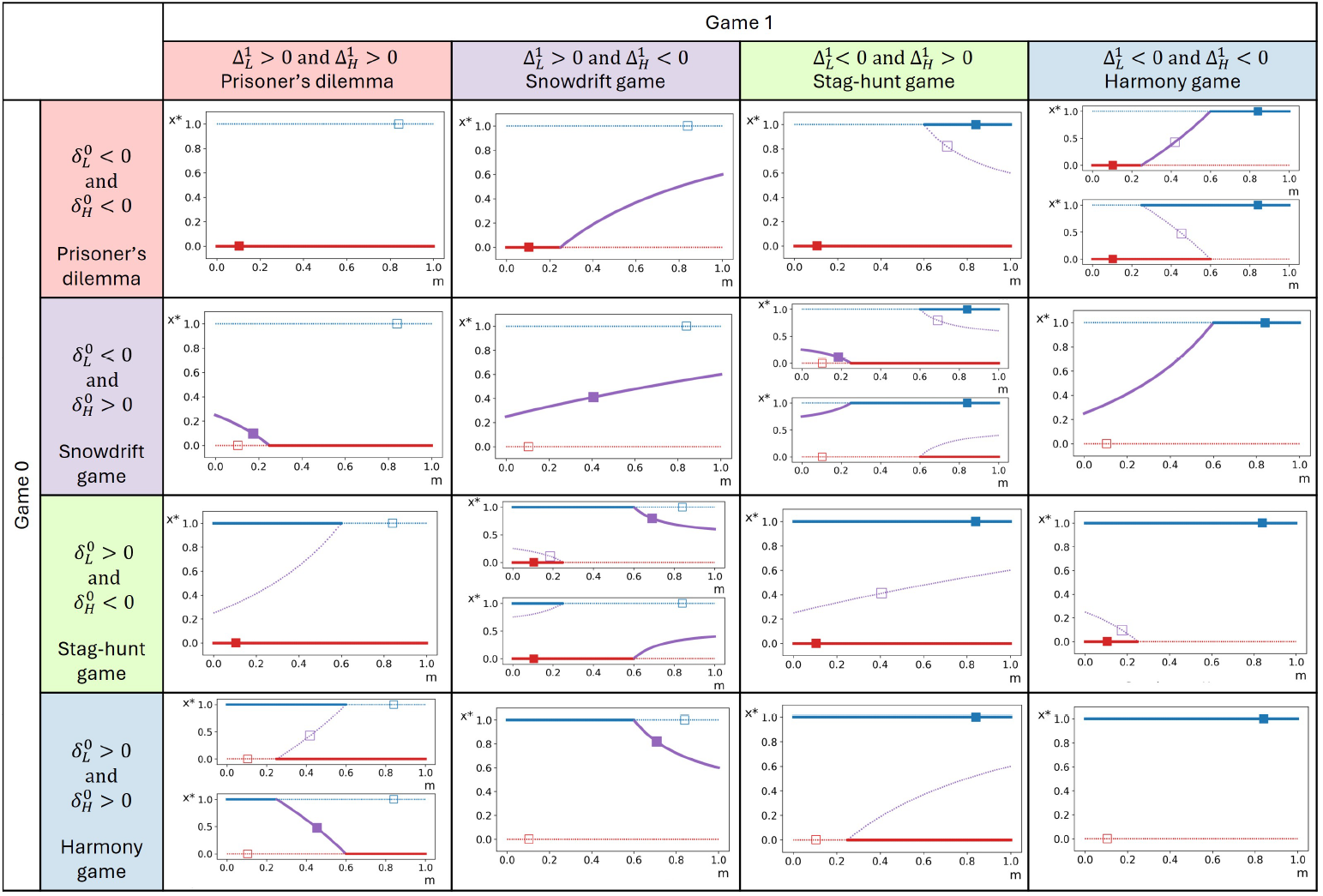
Bifurcation diagrams of the equilibrium cooperator frequency *x*\* as a function of environmental state *m* for all combinations of game types. In each panel, the horizontal axis represents *m* and the vertical axis represents the equilibria and their stability. The system admits three types of equilibria: *x*\* = 0 (red), *x*\* = 1 (blue), and an interior equilibrium *x*\* ∈ (0, 1) (purple), whose existence and stability depend on *m*. Solid lines denote stable equilibria and dotted lines unstable equilibria. Existence and stability depend on the combination of game types defined by Π(0) and Π(1). As in Fig. 2 of [11], rows correspond to the game structure of Π(0) (depleted environment) and columns correspond to the game structure of Π(1) (replete environment). Unless otherwise noted, parameter magnitudes are 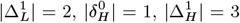, and 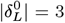. In four panels, a second bifurcation diagram is shown with 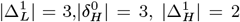, and 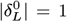 since in these cases the absolute values of the parameters qualitatively affect the bifurcation structure. Squares indicate the deterministic equilibria (*m, x*) from [20]: filled squares are stable and open squares unstable. result, individuals tend to stay at the behavior which is the most often optimal, across a range of environmental levels.

#### Stationary distributions of *m* and *x*

The bifurcation diagram of *x* as a function of *m* allows us, for most combinations of game types, to express the equilibrium fraction of cooperators as a function of the current environmental state. We denote this as *x*_*m*_. Substituting this relationship into the stochastic differential equation governing *m* yields a reduced description from which the stationary distribution of *m* can be derived, under the assumption of timescale separation.

As discussed above, we impose a reflecting boundary at *m* = 0. While the environmental state has no strict upper bound, the deterministic component of the dynamics stabilizes *m* between 0 and 1, and values *m >* 1 arise only due to stochastic fluctuations. The probability of *m* reaching very large values is therefore negligible. To obtain a stationary distribution on a finite domain, we impose an upper reflecting boundary at some sufficiently large value *m*_max_ ≫ 1 (for example, *m*_max_ = 10).

Under these assumptions, the stationary distribution of *m*, denoted *f* (*m*), is given by

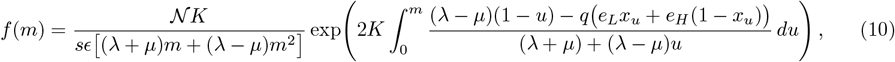

where𝒩is a normalization constant chosen such that 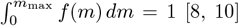. When the interior equilibrium 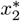 is stable, the stationary distribution of *x* can be obtained from *f* (*m*) by a change of variables (see Supplementary Information).

### 2.6 Numerical simulations

We also report results from two types of numerical simulations: simulations assuming type-scale separation (an SDE for *m* coupled to a deterministic function *x*_*m*_); and simulations without assuming timescale separation (an SDE for *m* coupled to an ODE for *x*).

## 3 Results

### 3.1 Behavior in regimes with timescale separation

We first consider regimes in which there is a clear separation of timescales between strategy evolution and environmental dynamics, namely *ϵ* → 0.

#### When long-term dynamics are a predictable mixture of behaviors

Because the effective game played by individuals depends on the state of the environment *m*, the evolution of behavior in the population also depends on the value of *m*. In a deterministic environment, resource levels converge to a stable equilibrium and the fraction of cooperators then evolve according to the payoff matrix associated with the equilibrium environmental state. When environmental dynamics are stochastic, however, the environmental state fluctuates around its equilibrium value, and the fraction of cooperators *x* can vary over time as a result.

As an illustrative example, consider a system in which Π(0) is a harmony game and Π(1) is a snowdrift game (Fig. 2a; game structure shown in the dilemma phase plane). In a deterministic environment, the only stable equilibrium of the eco-evolutionary system is the interior fixed point 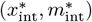 [20], indicated by a filled square in the bifurcation diagram, corresponding to coexistence of cooperators and defectors. In a noisy environment, by contrast, both full cooperation (*x* = 1) and the interior equilibrium can be stable, depending on the instantaneous value of *m* (see the bifurcation diagram). Consequently, the fraction of cooperators alternates between epochs of time near an intermediate value and epochs of time with full cooperation (see the trajectory of *x*).

The proportion of time the population spends in the fully cooperation state depends on payoff parameters and is reflected by a Dirac mass at *x* = 1 in the stationary distribution of *x*. If the game parameters extend the range of environmental states for which the effective game is a harmony game, the fraction of time that the population spends in the fully cooperative state also increases (see Figure 6 in Supplementary Information).

This example illustrates that stochasticity in the environment can qualitatively change the behavioral outcome in the population. In the absence of noise, the population always contains a mixture of cooperators and defectors. But in the presence of noise the population experiences epochs of time with a full cooperation, interspersed with epochs containing a mixture of cooperators and defectors. This overall pattern, although qualitatively different than the case without noise, is nonetheless somewhat intuitive: the changing environment moves between a harmony game (Π(0), where full cooperation would be expected in isolation) and a snowdrift game (Π(1), where a mixture of types would be expected in isolation), and the population composition also alternates between these two types of behavioral compositions.

Similar results hold for other pairs of games, as well—where the behavior with a noisy environment is distinct but comprehensible, given the behavior in each of the games Π(0) and Π(1) separately. For example, when Π(0) is a harmony game and Π(1) a prisoner’s dilemma, and when the effective game played for some intermediate value of *m* is a stag-hunt (Fig. 2b, dilemma phase portrait), the population will alternate between epochs of full cooperation and epochs of full defection (Fig. 2b). In the same setting with a noise-free environment the cooperator frequency can exhibit a stable limit cycle (oscillatory tragedy of the commons [27, 20]), oscillating between high and low frequencies of cooperation. In the presence of stochasticity we find similar behavior, although the oscillatory phases are noisy.

In summary, stochasticity in the environment can induce a qualitative change in the behavior of the population. The resulting pattern of behavior is sometimes intuitive to predict: as the environment varies the population adapts his behavior and has periods expressing behavior that is expected under the game in a poor environment (Π(0)), alternative with periods of behavior that is expected on the game induced by a rich environment (Π(0)).

#### When long-term behavior is not a predictable mixture of outcomes

For some combinations of type of games Π(0) and Π(1), the behavior of the population is not a simple or intuitive mixture of the behavior that might be expected for these two games in isolation.

For example, when Π(0) is a harmony game and Π(1) prisoner’s dilemma, and when the game for intermediate *m* is a snowdrift (namely, parameter regime 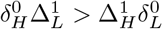), then the outcome is not simply a mixture of epochs associated with the two games. In isolation, a harmony game would lead to full cooperation, and a prisoner’s dilemma to full defection, but an interior equilibrium with a mixture of cooperators and defectors is stable for intermediate values of the environment *m* (Fig. 2c). The population will spend significant durations with a mixture of cooperators and defectors, even though this mixture would be unstable when *m* = 0 or *m* = 1. In sum, the range of behavior seen in such a population includes features that are not expected for either game Π(0) or game Π(1).

Environmental stochasticity can even turn one type of eco-evolutionary game into another type of game, over the long-term. When a stag-hunt is combined with another type of game, the eco-evolutionary behavior is no longer bistable, even though a stag-hunt game in isolation will produce bistability. Consider a prisoner’s dilemma and a stag-hunt game (Fig. 2d). In a noise-free environment there would be bistability [20]: some initial conditions will destine the population towards pure cooperation (*x* = 1, blue filled square in the bifurcation diagram). But in the presence of noise, once the environment reaches a state that leads to full defection (*x* = 0), the frequency of cooperators will subsequently be stuck at zero even as *m* changes, because *x* = 0 is stable for all the values of *m* (see trajectory of *x* in Figure 2d). This means that hysteresis caused by the noisy environment has fundamentally changed the nature of eco-evolutionary game: it has removed long-term characteristics of the stag-hunt game (namely, the possibility of pure cooperation) that would otherwise persist in the absence of environmental noise.

#### Long-term behavior for all combinations of all game types

Given the complexity across the examples discussed above, we seek a synthetic understanding of all possible dynamical outcomes in eco-evolutionary games with a noisy environment. We can provide such an account, at least in the regime *ϵ* → 0 with timescale separation, by considering all bifurcation diagrams for all pairs of game types, shown in Fig. 3. Across the 20 different qualitatively different regimes of games, we find that the observed range of behavioral composition in the population is typically larger in the presence of noise than in a deterministic environment.

For example, for a combination of a snowdrift game and a stag-hunt game, the population’s behavior includes several features previously described: it depends on the game played for intermediate values of *m* (prisoner’s dilemma or harmony game), the stag-hunt game can be effectively turned into a prisoner’s dilemma or into harmony game, and the behavior of the population over time is a mixture of behaviors typically associated with one or the other type of game.

We find a qualitative change of the behavior in noisy environments compared to deterministic environments in 14 of 20 cases. The qualitative behavior is qualitatively unchanged by noise, regardless of *K*, only if the equilibrium fraction of cooperators is stable in all states of the environment—that is, when Π(0) and Π(1) are the same types of games, when Π(0) is a stag-hunt game and Π(1) a prisoner’s dilemma, or when Π(0) is a harmony game and Π(1) a stag-hung game.

### 3.2 Impacts of carrying capacity and timescale of the environmental change

So far we restricted our analysis to cases when the stochasticity is sufficiently high and environmental dynamics slow compared to changes in behavior (small *K* and *ϵ*). In this section we investigate the impact of the size of the carrying capacity *K* (1*/K* determines the magnitude of noisy fluctuations in the environment) and of the timescale parameter *ϵ* on the stationary distributions of the environmental state and cooperator frequency. We first focus on system dynamics when one of the games is a snowdrift game.

In Fig. 4, Π(0) is a harmony game and Π(1) is a snowdrift game. In a deterministic environment (corresponding to *K*→∞), the only stable equilibrium is an interior equilibrium with a mixture of cooperators and defectors [20]. When the carrying capacity of environmental resource is large, the level of the environment remains close to this equilibrium value, which implies that there is almost always of mixture of cooperators and defectors *x* ∈ (0, 1) (see the distributions for *K* = 1000 in Fig. 4). But when the carrying capacity is smaller, the state of the environment is subject to stronger stochastical perturbation away from is deterministic equilibrium, including to environmental levels that induce qualitatively different effective games. Under timescale separation (*ϵ* →− 0), this implies that the fraction of cooperators moves away from its equilibrium in a noise-free environments (see the stationary distribution of *x* for *K* = 150 and *K* = 20 in Fig. 4a). In other words, in a noisy setting the behavior of the population is always a mix between harmony game and snowdrift game but the mix favors the equilibrium associated with a deterministic environment when the carrying capacity of resources is large, whereas it depends more on the game structure when the carrying capacity of the environmental resources is small.

**Figure 4:**
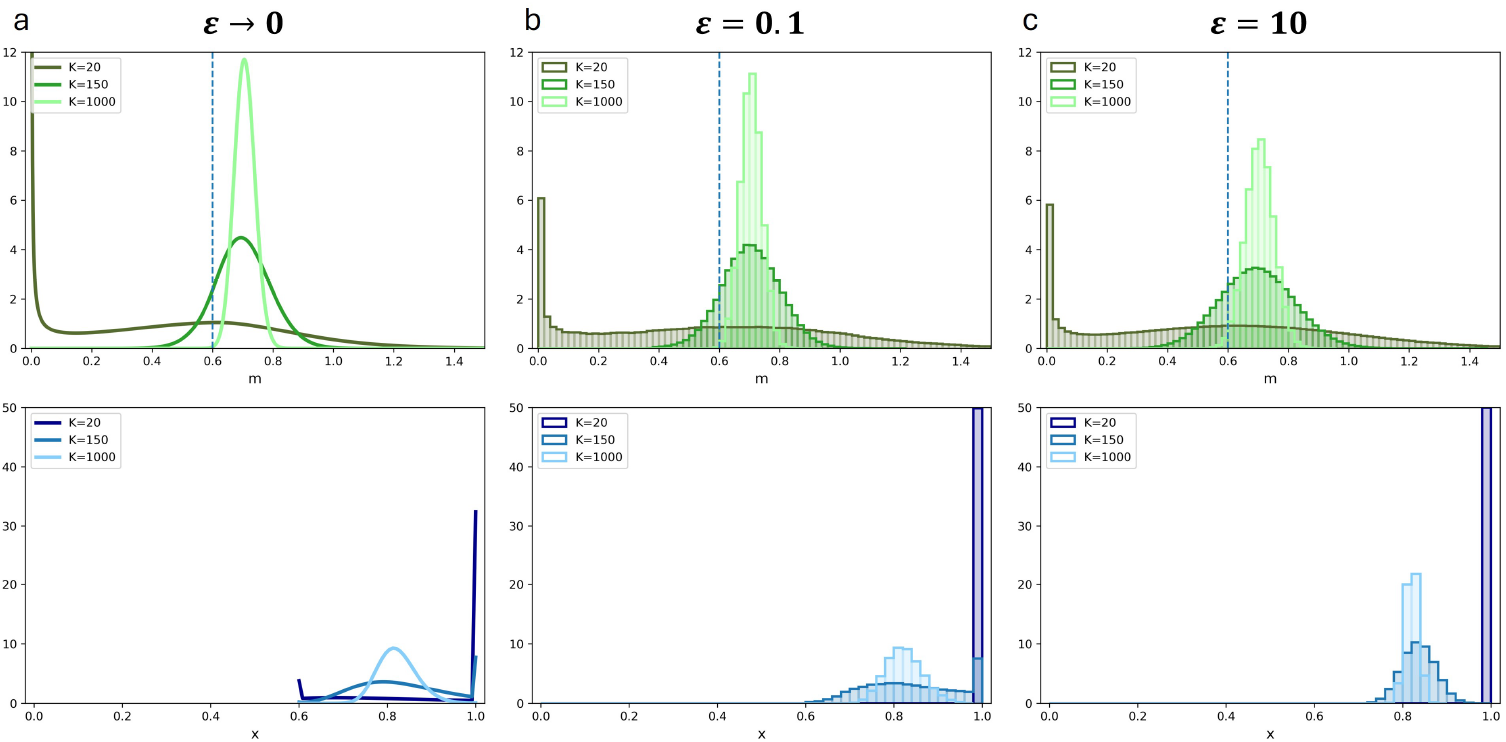
Impact of environmental carrying capacity *K* and timescale parameter *ϵ* on the stationary distribution of environment and cooperator frequency. Parameters are 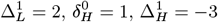, and 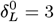, corresponding to a combination of harmony and snowdrift games (see dilemma phase plane in Fig. 2a). In each panel, the top row shows the stationary distribution of the environmental state *m*, and the bottom row shows the corresponding stationary distribution of the cooperator frequency *x*. The dotted vertical line (top panels) indicates the threshold value of *m* below which full cooperation is stable and above which an interior mixed equilibrium is stable (cf bifurcation diagram in Fig. 2a). Stationary distributions are shown for three values of *K* (shades of blue and green). For *K* = 1000 (weak noise), behavior is qualitatively the same as the deterministic case [20], with a stable mixed equilibrium. As *K* decreases, environmental fluctuations broaden and the population alternates between epochs of full cooperation and epochs with mixed strategies. In **a**, we take the limit *ϵ* → 0 and plot the analytical stationary distributions under strict timescale separation. In **b** and **c**, stationary distributions are obtained from simulations of the coupled SDE–ODE system at steady state.

When *ϵ* is large, *i*.*e*. when the dynamics of the environment are rapid and timescale separation does not hold, individuals have less time to equilibrate their strategies following an environmental perturbation. The level of cooperation alternates less often between a boundary equilibrium and an interior equilibrium. As a

If the carrying capacity is large enough, then the stationary distribution of *m* is concentrated around its equilibrium in a noise-free environment: *m* will spend most of its time at values for which the optimal strategy in the population agrees with the outcome in a noise-free environments (see *K* = 1000 and *K* = 150 in Fig. 4c).

Finally, as carrying capacity of the environment decreases, *m* is subject to greater stochasticity and it spends more time near 0 (see Fig. 4 for *K* = 20). This occurs because there is comparatively less noise for low values of *m*, so that *m* near 0 is “sticky”. As a result, for small *K* the behavior of the population is concentrated around the optimal behavior associated with whatever game corresponds to low resource level. Thus, for small carrying capacity and rapidly changing environments, the behavior of the population is determined by Π(0), and it can be qualitatively different from the one observed for large carrying capacity.

### 3.3 Noisy oscillations in the tragedy of the commons

In deterministic environments, [20] found that a stable limit cycle can arise for some pairs of games, provided the speed of environmental change is sufficiently low compared to the speed of strategic change (*ϵ < ϵ*_crit_, see Fig. 5a and b). This cyclical dynamic decreases the mean fitness of the population, hence the term “oscillating tragedy of the commons” (Fig 5a, b and c, [27]). However, if the speed of environmental change is closer to that of behavioral change (*ϵ > ϵ*_crit_), the oscillatory tragedy of the commons is averted and replaced by a stable mixture of cooperators and defectors (Fig. 5c).

**Figure 5:**
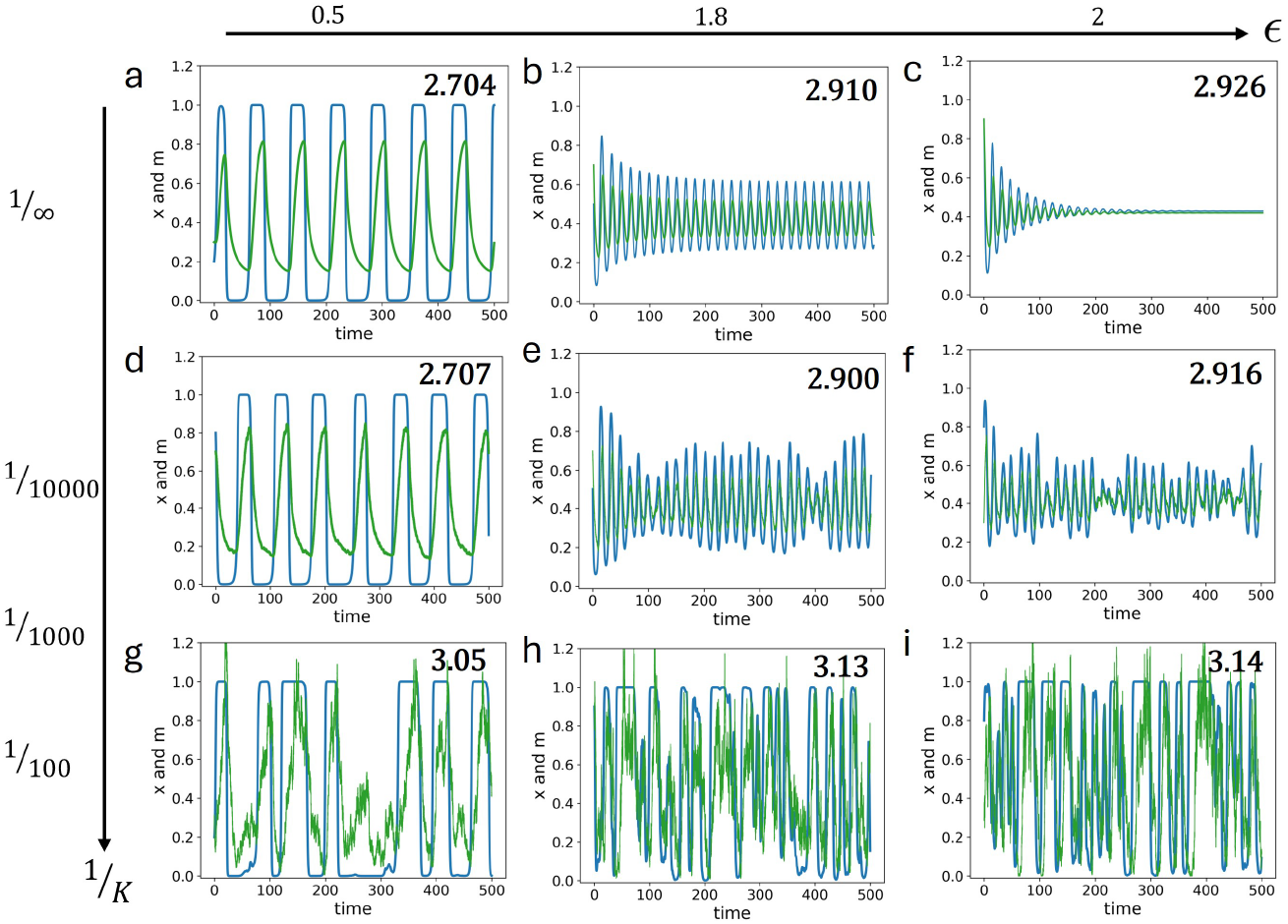
A noisy, oscillating tragedy of the commons. Parameters (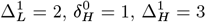, and 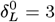) are chosen so that there is a harmony game in the rich state, prisoner’s dilemma in the poor state, and a stag-hunt game for intermediate values of *m* (see dilemma phase plane in Fig. 2b). The green curves show the trajectory of environmental state *m*, and the blue curves the corresponding cooperator frequency *x*. Each panel (upper right) reports the population mean fitness. The top row shows the deterministic limit (*K* → ∞), where oscillations occur only for *ϵ < ϵ*_crit_ ≈ 1.88 [20]. In the presence of noise (*K <* ∞, rows), oscillations persist regardless of timescale parameter *ϵ* (columns).

We find similar behavior in the presence of noise—except for one notable qualitative difference. When the resource has a finite carrying capacity, environmental state is noisy and the fraction of cooperators oscillates stochastically—much like a noisy version of the oscillating tragedy of the commons (Fig. 5d, e, f, g, h, i). Importantly, these stochastic oscillations persist regardless of the environmental timescale, *ϵ*. When environmental dynamics are noisy and rapid, the oscillations in cooperation have a lower magnitude and this partly rescues mean fitness; but a population in a noisy environment cannot escape the oscillating tragedy of the commons.

## 4 Discussion

Our central result is that environmental stochasticity can qualitatively alter eco-evolutionary outcomes. When environmental resources fluctuate due to noise, long-term behavior is no longer determined solely by the deterministic equilibrium of the coupled system; nor must it be a simple mixture of the behavior in a rich or poor environments. As the environmental carrying capacity decreases and fluctuations grow, this mixture can expand the range of observed behaviors, eliminate bistability that would otherwise persist, and in some cases force the dominance of a single strategy. Thus, even when the underlying games and feedback structure are unchanged, stochastic environmental dynamics can radically reshape both behavioral composition and resource levels.

Our study adds to a growing body of work showing that stochasticity can alter evolutionary outcomes. Demographic noise in a finite population of agents [24, 5], fluctuations in payoff observations [22], and variability in social environments [23] can all qualitatively modify the evolution of cooperation. Likewise, environmental variability has been shown to facilitate fixation in models without feedback between behavior and environmental state [1, 2]. Our study differs in an essential respect: we incorporate demographic stochasticity directly into the environmental resource within a coupled eco-evolutionary framework. In this setting, noise does not act on payoffs or strategy updating directly; instead, it perturbs the environmental state, which in turn reshapes the strategic incentives faced by all individuals.

This distinction has important consequences. Previous studies have shown that asymmetric payoff noise can effectively transform one game into another [23]. Here, by contrast, game transformation arises endogenously through stochastic environmental feedbacks that affect cooperators and defectors symmetrically. Even though the payoff matrices Π(0) and Π(1) are fixed, environmental fluctuations can shift the effective game experienced by the population and thereby alter the qualitative long-run outcome.

Our results also clarify how stochastic eco-evolutionary dynamics relate to the deterministic framework developed in [20]. In the deterministic limit of large environmental carrying capacity, environmental trajectories converge to fixed points or limit cycles, and long-run behavior is governed by the equilibrium environmental state. When environmental fluctuations are introduced, long-run behavior instead reflects a weighted mixture of the strategies favored under Π(0) and Π(1), with the weights determined by the stationary distribution of environmental states. As environmental carrying capacity decreases, this distribution broadens, and the range of observed behaviors expands accordingly. In this sense, our analysis extends the deterministic theory to finite-resource systems in which environmental noise cannot be neglected.

The relative timescale of environmental and strategic change remains a central feature that determines longterm patterns and even our analytical approach. As in the deterministic model, slow environmental dynamics can generate oscillatory tragedies of the commons. However, in the presence of stochasticity, these oscillations persist even outside the deterministic parameter regime that supports stable limit cycles. When environmental change is rapid and noisy, individuals cannot fully track instantaneous environmental states; instead, behavior tends toward the strategy that is most often optimal across fluctuating conditions. In the case of an oscillatory tragedy of the commons, stochastic fluctuations amplify the tendency toward fixation at the boundaries, thereby sustaining noisy cycles even when deterministic dynamics would produce convergence to a stable mixed equilibrium.

The framework developed here applies broadly to biological and social systems in which strategic behavior feeds back on a fluctuating environment. In particular, it provides a foundation for studying resource management problems under demographic uncertainty, such as optimal harvesting of renewable stocks [13, 16], where environmental variability is intrinsic rather than exogenous.

Nonetheless, our results have been derived under a range of simplifying assumptions that might be relaxed in future work. Our analytical approaches rely on strict separation of timescales between environmental and strategic dynamics. Outside this regime, we characterized long-term behavior numerically. A full analytical treatment of the general case would require analysis of two coupled stochastic differential equations. Developing such a theory remains an important analytical challenge.

We have focused on a renewing resource whose intrinsic dynamics are governed by a birth–death process. This represents one specific class of environmental stochasticity. In contrast, [20] also considered environments characterized by deterministic decay or tipping-point dynamics. Extending the present analysis to incorporate stochasticity in these alternative forms of environmental dynamics would clarify how general our conclusions are across different ecological regimes.

In addition, we have assumed that individuals affect the environment deterministically, so that stochasticity arises solely from intrinsic birth and death events in the resource. In many settings, however, harvesting itself may be partly stochastic, introducing an explicit dependence on *x* in the noise term of the environmental dynamics. Such contributions could be comparable in magnitude to intrinsic demographic fluctuations and may generate additional qualitative effects. Understanding how behavioral stochasticity interacts with environmental feedback is therefore a natural extension of the present model.

Several structural extensions of the game-theoretic framework are also possible. We have focused on linear two-player games that collectively influence a shared environment. The model could be generalized to multiplayer public goods games [17], asymmetric interactions [9], or nonlinear payoff structures [25]. Likewise, individuals could choose from a continuous action space representing harvesting effort, and environmental dynamics could be spatially extended or heterogeneous, allowing for spatial variation in behavior and migration across local environmental states [14, 28]. Each of these extensions would enrich the interaction between stochasticity and eco-evolutionary feedback.

Finally, our analysis has focused on myopic agents who respond to current payoffs. An important open question is how forward-looking behavior interacts with environmental stochasticity. In deterministic eco-evolutionary models, forecasting agents can coexist with myopic agents and increase population mean payoff [21]. When the environment is noisy, however, forecasting becomes a problem of inference under uncertainty. Whether predictive strategies, institutional forecasting, or incentive mechanisms such as subsidies or punishment [3, 4, 26] can stabilize cooperation in stochastic eco-evolutionary systems remains an important direction for future research.

## Code availability

The script for performing simulations and plotting the figures is available through the following link : https://github.com/fantine-bodin/Eco-evo_games_noisy_env.

## Statement of contributions

Fantine Bodin, Guocheng Wang and Joshua B. Plotkin designed the study. The model for strategy evolution and for environmental feedbacks is taken from the literature. Fantine Bodin, Joshua B. Plotkin and Guocheng Wang developed a discrete model for noisy environmental dynamics and derived its SDE limit. Fantine Bodin performed the mathematical analyses and the simulations and wrote a first version of the manuscript. Fantine Bodin, Guocheng Wang and Joshua B. Plotkin contributed to the revisions.

## Acknowledgments

The authors acknowledge members of the Plotkin lab at the University of Pennsylvania and members of Princeton Theory Tea for helpful discussions.

## References

[1] Peter Ashcroft, Philipp M. Altrock, and Tobias Galla. Fixation in finite populations evolving in fluctuating environments. 11(100):20140663.

[2] Michael Assaf, Mauro Mobilia, and Elijah Roberts. Cooperation dilemma in finite populations under fluctuating environments. 111(23):238101.

[3] Xiaojie Chen and Attila Szolnoki. Punishment and inspection for governing the commons in a feedback-evolving game. 14(7):e1006347.

[4] Hiroaki Chiba-Okabe and Joshua B. Plotkin. Can institutions foster cooperation by wealth redistribution? 21(212):20230698.

[5] George W. A. Constable, Tim Rogers, Alan J. McKane, and Corina E. Tarnita. Demographic noise can reverse the direction of deterministic selection. 113(32).

[6] Peter Czuppon and Arne Traulsen. Understanding evolutionary and ecological dynamics using a continuum limit.

[7] Sylvie Estrela, Eric Libby, Jeremy Van Cleve, Florence Débarre, Maxime Deforet, William R. Harcombe, Jorge Peña, Sam P. Brown, and Michael E. Hochberg. Environmentally mediated social dilemmas. 34(1):6–18.

[8] Crispin W. Gardiner. Stochastic methods: a handbook for the natural and social sciences. Number 13 in Springer series in synergetics. Springer, 4th ed edition.

[9] Christoph Hauert, Camille Saade, and Alex McAvoy. Asymmetric evolutionary games with environmental feedback. 462:347–360.

[10] Werner Horsthemke. Noise-Induced Transitions: Theory and Applications in Physics, Chemistry, and Biology. Number v.15 in Springer Series in Synergetics Ser. Springer Berlin/Heidelberg.

[11] Hiromu Ito and Masato Yamamichi. A complete classification of evolutionary games with environmental feedback. 3(11):pgae455.

[12] Taylor A. Kessinger, Corina E. Tarnita, and Joshua B. Plotkin. Evolution of norms for judging social behavior. 120(24):e2219480120.

[13] Sarah B M Kraak. Exploring the ‘public goods game’ model to overcome the tragedy of the commons in fisheries management. 12(1):18–33.

[14] Yu-Hui Lin and Joshua S. Weitz. Spatial interactions and oscillatory tragedies of the commons. 122(14):148102.

[15] Martin A. Nowak and Karl Sigmund. Evolutionary dynamics of biological games. 303(5659):793–799.

[16] Thomas W. Schoener. The newest synthesis: Understanding the interplay of evolutionary and ecological dynamics. 331(6016):426–429.

[17] Yanxuan Shao, Xin Wang, and Feng Fu. Evolutionary dynamics of group cooperation with asymmetrical environmental feedback. 126(4):40005.

[18] Christine Taylor, Drew Fudenberg, Akira Sasaki, and Martin A Nowak. Evolutionary game dynamics in finite populations. Bulletin of mathematical biology, 66(6):1621–1644, 2004.

[19] Peter D. Taylor and Leo B. Jonker. Evolutionary stable strategies and game dynamics. 40(1):145–156.

[20] Andrew R. Tilman, Joshua B. Plotkin, and Erol Akçay. Evolutionary games with environmental feedbacks. 11(1):915.

[21] Andrew R Tilman, Vítor V Vasconcelos, Erol Akçay, and Joshua B Plotkin. The evolution of forecasting for decision-making in dynamic environments. 2(4):26339137231221726.

[22] Guocheng Wang, Qi Su, Long Wang, and Joshua B. Plotkin. The evolution of social behaviors and risk preferences in settings with uncertainty. 121(30):e2406993121.

[23] Guocheng Wang, Qi Su, Long Wang, and Joshua B. Plotkin. Evolution of social behaviors in noisy environments. Version Number: 1.

[24] Guocheng Wang, Qi Su, Long Wang, and Joshua B. Plotkin. Reproductive variance can drive behavioral dynamics. 120(12):e2216218120.

[25] Xin Wang and Feng Fu. Eco-evolutionary dynamics with environmental feedback: Cooperation in a changing world. 132(1):10001.

[26] Xin Wang, Zhiming Zheng, and Feng Fu. Steering eco-evolutionary game dynamics with manifold control. 476(2233):20190643.

[27] Joshua S. Weitz, Ceyhun Eksin, Keith Paarporn, Sam P. Brown, and William C. Ratcliff. An oscillating tragedy of the commons in replicator dynamics with game-environment feedback. 113(47).

[28] Tianyong Yao and Daniel B. Cooney. Spatial pattern formation in eco-evolutionary games with environment-driven motion. Version Number: 1.

