## Supplementary Information for "Eco-evolutionary games in noisy environments"

### Eco-evolutionary games in noisy environments: Supplementary Information

May 20, 2026

[1]Department of Biology, University of Pennsylvania, Philadelphia, PA 19104
[2]Institut de Biologie de l'ENS (IBENS), École Normale Supérieure - PSL, 46 rue d'Ulm, 75005 Paris, France
[3]Center for Mathematical Biology, University of Pennsylvania, Philadelphia, PA 19014

Contact:

### 1 Supplementary Information

#### 1.1 List of variables

| Variable | Definition | Typical values |
|---|---|---|
| $\delta_H^0$ | $S_0 - P_0$, incentive to switch to low-impact strategy when all others have high-impact strategy in a poor environment, *i.e.* incentive to lead an environmental movement | $[\![-4;4]\!]$ |
| $\Delta_H^1$ | $P_1 - S_1$, incentive to switch to high-impact strategy when all others have high-impact strategy in a rich environment, *i.e.* incentive to follow the gold rush | $[\![-4;4]\!]$ |
| $\delta_L^0$ | $R_0 - T_0$, incentive to switch to low-impact strategy when all others have low-impact strategy in a poor environment, *i.e.* incentive to follow an environmental movement | $[\![-4;4]\!]$ |
| $\Delta_L^1$ | $T_1 - R_1$, incentive to switch to high-impact strategy when all others have low-impact strategy in a rich environment, *i.e.* incentive to lead the gold rush | $[\![-4;4]\!]$ |
| $N$ | Size of the population of cooperators and defectors | 1000 |
| $x$ | Proportion of cooperators in the population | $[0,1]$ |
| $s$ | Selection coefficient | 0.1 |
| $M$ | Number of environmental resources | $[0,+\infty)$ |
| $K$ | Carrying capacity of environmental resources | $[10,+\infty)$ |
| $m$ | Level of environmental resources rescaled by $K$ | $[0,+\infty)$ |
| $\lambda$ | Birth rate of environmental resources | 1.2 |
| $\mu$ | Death rate of environmental resources | 0.7 |
| $e_H$ | Harvest effort of high-impact individuals | 0.56 |
| $e_L$ | Harvest effort of low-impact individuals | 0.1 |
| $q$ | Parameter which maps degradation pressure by the individuals to the rate of reduction in the resource | 0.8 |
| $\epsilon$ | Rate of environmental changes compared to the rate of strategies updating | $[0,10]$ |

**Table 1:** List of variables used in the model and values used for the figures and for the simulations

#### 1.2 Model of joint eco-evolutionary dynamics

As described in the main text, we assume a discrete-state model for strategy evolution and environmental dynamics. The level of cooperation $x$ and the level of environmental resources $m$ follow a discrete Markov Chain process in continuous time with the following transition rates:

- $x$ increases from $x$ to $x + 1/N$ at rate $(1-x)x\phi(\pi_H(x,m), \pi_L(x,m))$
- $x$ decreases from $x$ to $x - 1/N$ at rate $x(1-x)\phi(\pi_H(x,m), \pi_L(x,m))$
- $m$ increases from $m$ to $m + \frac{1}{K}$ at rate $s\epsilon\left[\lambda m - \frac{q}{2}m(e_L x + e_H(1-x))\right]$
- $m$ decreases from $m$ to $m - \frac{1}{K}$ at rate $s\epsilon\left[\mu m + (\lambda-\mu)m^2 + \frac{q}{2}m(e_L x + e_H(1-x))\right]$

#### 1.3 Derivation of associated SDEs

When the size of the population $N$ and the carrying capacity of the environment $K$ are sufficiently large, we can approximate the discrete eco-evolutionary model using a continuous limit.

We first focus on deriving an ordinary differential equation (ODE) for the proportion of cooperators, when $N$ is large.

$$\mathbb{E}(\Delta x) = \frac{1}{N}(1-x)x\phi(\pi_H, \pi_L) - \frac{1}{N}x(1-x)\phi(\pi_L, \pi_H) \tag{1}$$

Given $s \ll 1$, we perform a Taylor expansion of $\phi(\pi_i, \pi_j) = 1/(1+\exp[s(\pi_i - \pi_j)]$ and obtain after rescaling time by $N$, setting $\Delta t = 1/N$ and letting $N \to \infty$:

$$\dot{x} = \mathbb{E}(\Delta x)/(1/N) = sx(1-x)(\pi_L - \pi_H) \tag{2}$$

We can derive a continuous model for the environment dynamics under the limit of $K$ large, without neglecting environmental demographic noise. We obtain a stochastic differential equation (SDE) for $m$ (Czuppon et al. 2020) after rescaling time by $K$, setting $\Delta t = 1/K$ and rescaling $\epsilon$ by $K/N$:

$$\begin{aligned}
\mathbb{E}(\Delta m) &= \frac{1}{K}s\epsilon\left[\lambda m - \frac{q}{2}m(e_L x + e_H(1-x))\right] - \frac{1}{K}s\epsilon\left[\mu m + (\lambda-\mu)m^2 + \frac{q}{2}m(e_L x + e_H(1-x))\right] \\
\mathbb{E}(\Delta m^2) &= \frac{1}{K^2}s\epsilon\left[\lambda m - \frac{q}{2}m(e_L x + e_H(1-x))\right] + \frac{1}{K^2}s\epsilon\left[\mu m + (\lambda-\mu)m^2 + \frac{q}{2}m(e_L x + e_H(1-x))\right] \\
dm &= \mathbb{E}(\Delta m)/(1/K)\,dt + \sqrt{\mathbb{E}(\Delta m^2)/(1/K)}\,dW_t \\
&= s\epsilon[(\lambda-\mu)m(1-m) - qm(e_L x + e_H(1-x))]\,dt + \sqrt{s\epsilon\frac{(\lambda+\mu)m + (\lambda-\mu)m^2}{K}}\,dW_t
\end{aligned} \tag{3}$$

with $dW_t$ the Brownian motion.

#### 1.4 Regime of timescale separation

##### Existence and stability of equilibria of $x$ as a function of $m$: bifurcation diagrams

If $\epsilon$ is sufficiently small, the environment is changing slowly compared to the evolution of strategies in the population. This means that at each time the environment is changing, the population has time to adapt its behavior to equilibrium before the environment changes again (timescale separation). The ordinary differential equation that governs $x$ can then be written as:

$$\begin{aligned}
\dot{x} &= sx(1-x)(\pi_L(x,m) - \pi_H(x,m)) \\
&= sx(1-x)((1-m)(R_0 x + S_0(1-x)) + m(R_1 x + S_1(1-x)) - (1-m)(T_0 x + P_0(1-x)) + m(T_1 x + P_1(1-x))) \\
&= sx(1-x)(a(m)x + b(m)) \\
&\text{with } a(m) = (\Delta_H^1 - \Delta_L^1 - \delta_L^0 + \delta_H^0)m + (\delta_L^0 - \delta_H^0) \text{ and } b(m) = (-\delta_H^0 - \Delta_H^1)m + \delta_H^0
\end{aligned} \tag{4}$$

The fraction of cooperators $x$ has three possible equilibria:

$$\left\{x_0^* = 0, x_1^* = 1, x_2^* = -\frac{b(m)}{a(m)}\right\} \tag{5}$$

To analyze their stability, we calculate the second derivative of $x$ at these points:

$$\frac{d}{dx}\frac{dx}{dt}\bigg|_{x=x_0^*} = sb(m), \ \frac{d}{dx}\frac{dx}{dt}\bigg|_{x=x_1^*} = -s(a(m)+b(m)), \ \frac{d}{dx}\frac{dx}{dt}\bigg|_{x=x_2^*} = -s\frac{b(m)}{a(m)}(a(m)+b(m)) \tag{6}$$

The equilibrium $x_2^*$ is feasible only if it lies between 0 and 1. Note that $x_2^* \in [0,1] \Leftrightarrow$ ($x_0^*$ and $x_1^*$ stable) or ($x_0^*$ and $x_1^*$ unstable). Thus, $x_2^* \in [0,1]$ and stable $\Leftrightarrow$ $x_0^*$ and $x_1^*$ unstable.

All that remains is to study the stability of $x_0^*$ and $x_1^*$.

$$\begin{aligned} x_0^* \text{ stable } &\Leftrightarrow sb(m) < 0 \Leftrightarrow (-\delta_H^0 - \Delta_H^1)m + \delta_H^0 < 0 \\ x_1^* \text{ stable } &\Leftrightarrow -s(a(m)+b(m)) < 0 \Leftrightarrow (-\delta_L^0 - \Delta_L^1)m + \delta_L^0 > 0 \end{aligned} \tag{7}$$

If $\Delta_H^1$ and $\delta_0^H$ (resp. $\Delta_L^1$ and $\delta_L^0$) have opposite signs, $x_0^*$ (resp. $x_1^*$) is either stable or unstable for all the values of $m \in [0,1]$. If they have the same sign, $x_0^*$ (resp. $x_1^*$) switches at value $m_H = \dfrac{\delta_H^0}{\delta_H^0 + \Delta_H^1} \in (0,1)$ (resp. $m_L = \dfrac{\delta_L^0}{\delta_L^0 + \Delta_L^1}$) between stable and unstable or between unstable and stable.

Using these stability conditions we can plot a bifurcation diagram of the three equilibria of $x^*$ as a function of $m$, for each combination of $\delta_L^0, \delta_H^0, \Delta_H^1, \Delta_L^1$ (Fig. **??**).

##### Stationary distribution of $m$ and $x$

The bifurcation diagram of $x$ as a function of $m$ allows us to write $x$ as a function of $m$ (denoted $x_m$) for most combinations of game types. This expression can be substituted into the SDE that describes the behavior of $m$. Thus, we can find the stationary distribution of $m$ (assuming timescale separation).

As described in the main text, we assume that 0 is a reflecting boundary for $m$. The level of environmental resources has no upper boundary. However, the deterministic dynamics of $m$ stabilizes it between 0 and 1 and $m$ can only exceed 1 due to the stochastic component. The probability that $m$ goes to large values is quickly negligible. To obtain a stationary distribution on a finite interval, we therefore assume that $m$ has an upper reflecting boundary at some value $m_{\max} \gg 1$, for example at $m_{\max} = 10$. The stationary distribution of $m$, denoted by $f$, is then

$$f(m) = \frac{\mathcal{N}K}{s\epsilon[(\lambda+\mu)m + (\lambda-\mu)m^2]}\exp\left(2\int_0^m \frac{(\lambda-\mu)(1-u) - q(e_L x_u + e_H(1-x_u))}{(\lambda+\mu)+(\lambda-\mu)u} K\,du\right) \tag{8}$$

with $\mathcal{N}$ a constant so that $\int_0^{m_{\max}} f(m)dm = 1$ (Gardiner 2009; Horsthemke 2006).

When $x_2^*$ is stable, we can deduce the stationary distribution of $x$, denoted by $g$, thanks to a change of variable:

$$g(x) = f(m(x))\left|\frac{dm}{dx}\right| \tag{9}$$

For $x = 0$ (resp. $x = 1$), $g(x)$ is the integral of $f(m)$ over all the interval of $m$ for which $x = 0$ (resp. $x = 1$) is stable. For $x = \dfrac{-b(1)}{a(1)}$, $g(x) = \int_1^{+\infty} f(m)$. The stationary distribution of $x$ describes the proportion of time that the population spends at each level of cooperation.

The analytic expression of $f$ and $g$ are integrated numerically in Python with the library SciPy for producing figures in the main text. To illustrate the trajectory of $x$ under timescale separation, and to check our analytical stationary distributions, we performed simulations with Python of the SDE for $m$, substituting $x_m$. The analytical stationary distributions are mapped to the histogram obtained from simulations by taking into account the size of the bins.

#### 1.5 Stationary distributions without timescale separation

To determine the impact of $K$ and $\epsilon$ on the behavior of the population, we simulated the associated SDEs over a sufficiently long time to obtain good approximations of the stationary distributions for $m$ and $x$. Simulations of the SDEs were performed in Python, using the Euler method.

#### 1.6 Supplementary Figure

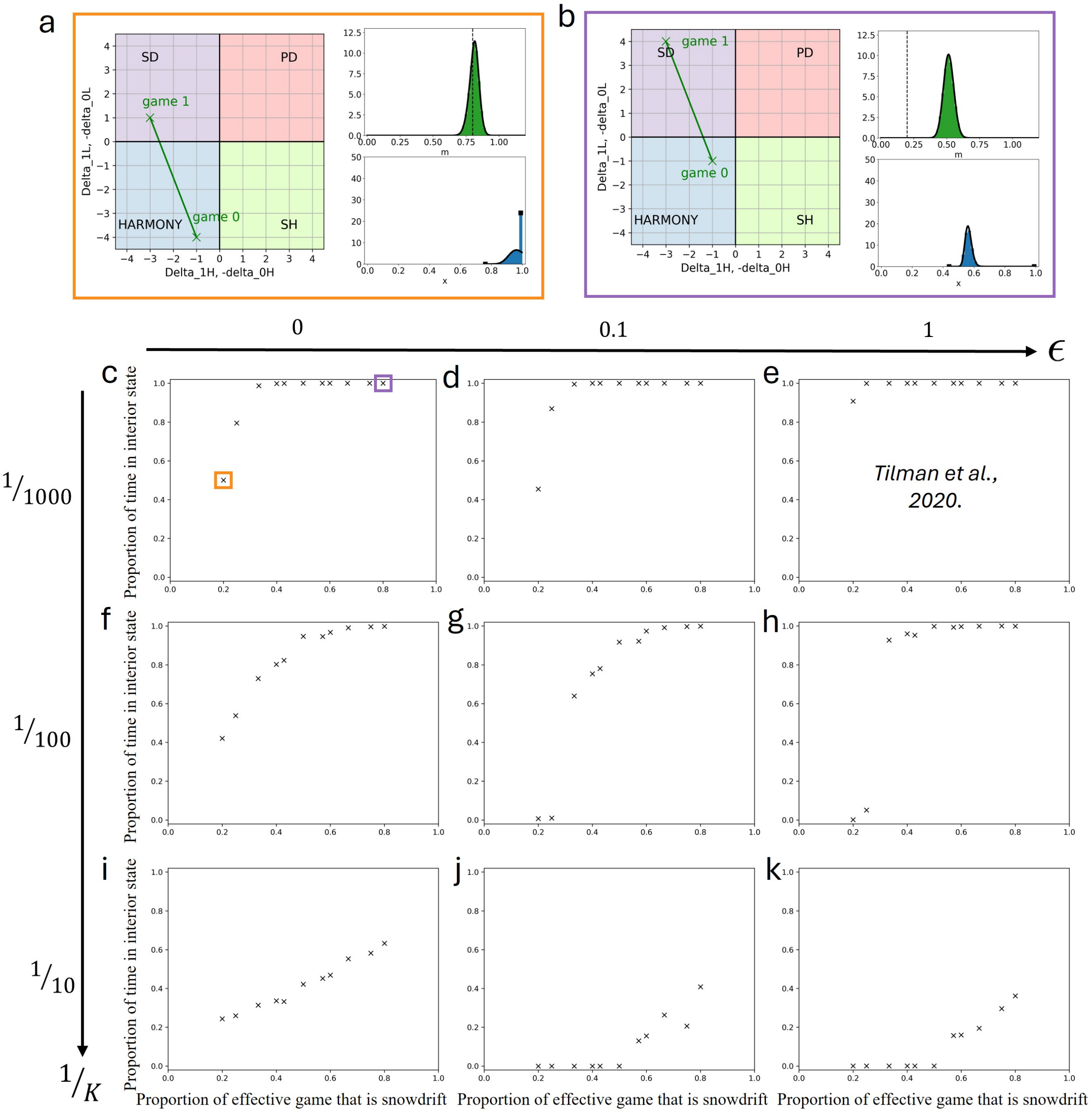


**Figure 1: Impact of the proportion of effective games that are snowdrift on cooperator frequency $x$, for different values of carrying capacity $K$ and timescale $\epsilon$.** Here $\Pi(0)$ is a harmony game and $\Pi(1)$ is a snowdrift game (see dilemma phase plane of Fig. **??**a.). **a, b.** Dilemma phase plane and stationary distributions of $m$ (in green, on the top) and $x$ (in blue, on the bottom) for two different proportions of effective game that is snowdrift (it corresponds to the proportion of the green line in the snowdrift quadrant on the dilemma phase plane). **a.** The proportion of effective game that is snowdrift is 0.2. According to the stationary distribution, the level of cooperation $x$ is half of the time at 1 and half of the time at an intermediate value between 0 and 1. **b.** The proportion of effective game that is snowdrift is 0.8. According to the stationary distribution, the level of cooperation $x$ is always at an intermediate value and never at 0 and 1. **c to k.** For each panel, the x-axis represents the proportion of effective game which is a snowdrift game. The y-axis represents the proportion of time that $x$ spends at an internal value between 0 and 1, *i.e.* not at the boundaries. Each cross is the result of a simulation ran with a certain couple $(\delta_L^0, \Delta_L^1)$ which determines the proportion of effective game which is a snowdrift game. **c, d, e.** $K = 1000$. **f, g, h.** $K = 100$. **i, j, k.** $K = 10$. **c, f, i.** Simulations with strict timescale separation, where $\epsilon$ is considered sufficiently low so that $x$ is always at equilibrium. **d, g, j.** $\epsilon = 0.1$. **e, h, k.** $\epsilon = 1$. In panel i, $K$ and $\epsilon$ are sufficiently high so that whatever is the proportion of game that is snowdrift, we obtain a behavior similar as the behavior without noise described by Tilman et al. 2020.
